# Motif Scraper: a Cross-Platform, Open-Source Tool for Identifying Degenerate Nucleotide Motif Matches in FASTA Files

**DOI:** 10.1101/265108

**Authors:** Elisha D. O. Roberson

## Abstract

**Summary:** Many genomic features are defined not by exact sequence matches, but by degenerate nucleotide motifs that represent multiple compatible matches. While there are databases cataloging genomic features, such as the location of transcription factor motifs, for commonly used model species, identifying the locations of novel motifs, known motifs in non-model genomes, or known motifs in personal whole-genomes is difficult. I designed motif scraper to overcome this limitation, allowing for efficient, multiprocessor motif searches in any FASTA file.

**Availability and implementation:** The motif scraper package is available via PyPI, and the Python source is available on GitHub at https://github.com/RobersonLab/motif_scraper.

## A. Introduction

Genomic features can often be described by sequence motifs, rather than exact sequence matches. Particularly important examples of this property are proximal promoter elements that bind transcription factors, and proteins that bind at enhancers and insulators. In these cases, the binding protein does not find an exact sequence match, but rather binds a range of sequences with compatible charge profiles for the protein binding interface. Using methods such as ChIP-Seq, the binding sequences for these factors can be determined and represented as a sequence motif using IUPAC-approved degenerate nucleotide codes. Other important features are exact matches, such as the match between a microRNA (**miR**) and seed sequences in the 3' untranslated region (**UTR**) of a targeted gene. Many databases exist cataloging the location of transcription factor motifs (Kaplun, et al., 2016; Kel, et al., 2003; Knuppel, et al., 1994; Matys, et al., 2006; Wingender, 1988; Wingender, 2008; Wingender, et al., 1996), miRNA binding sites (Andres-Leon, et al., 2015; Dweep, et al., 2014; Griffiths-Jones, 2004; Griffiths-Jones, et al., 2006; Griffiths-Jones, et al., 2008; Kozomara and Griffiths-Jones, 2014; Lagana, et al., 2012; Prabahar and Natarajan, 2017), and genome-editing sites (Gratz, et al., 2014; Heigwer, et al., 2014; Liu, et al., 2015; Montague, et al., 2014; Naito, et al., 2015; Stemmer, et al., 2015; Xiao, et al., 2014). However, these databases are often restricted to commonly used model species. Newly sequenced species are likely never to be included, and model species may lag behind the release of new genome drafts. Furthermore, many individual, phased whole genomes are being generated. The databases of sequence motifs are designed relative to a reference sequence, rather than to personal genomes. Inspired by previous work to identify a specific subset of CRISPR/Cas9 sites (Roberson, 2015), my goal for motif scraper was to instead develop a more general purpose motif searching tool that would have broader use. Motif scraper fills this annotation gap by allowing for the specification degenerate sequence motifs and reporting the position and composition of all matches in a FASTA file, which could be a personal genome, a reference genome, or a set of genomic slices, such as all the 3' UTRs of protein coding genes.

## B. Methods

Motif scraper was designed in Python, and is compatible with both Python 2 and 3. The ability to read FASTA formatted files and generate FASTA indexes is provided by pyfaidx (Shirley, et al., 2015). Motifs are specified as a text string with using IUPAC degenerate bases, which are converted internally into a regular expression and compiled by the regex package. This allows for detection of overlapping motifs. One or more specific regions or a specific strand relative to the reference can be specified for targeted search. By default all contigs in the FASTA file are searched for both + and − strands. The multiprocessor Python package handles the use of multiple computer cores, searching each target region / strand separately. Each hit is reported with the contig, start position, end position, strand, sequence, and matching motif in the output file. The code is available under an MIT license, stored on GitHub, and distributed through the Python Package Index (PyPI). Compatibility with Python 2 and 3 is assessed with every repository commit using Travis CI service.

## C. Results

### C.1. Identification of mock transcription factor binding motifs

As a benchmark, I calculated a faux consensus sequence for two DNA binding proteins: CCAAT/Enhancer Binding Protein Beta (**CEBPB**) and CCCTC-Binding Factor (**CTCF**). I downloaded their Position Weight Matrices for *Homo sapiens* from Jaspar (Mathelier, et al., 2016). I then calculated the fraction of weight at each position attributable to each base. At each position I considered a base contributing at least 5% of overall weight to be a possible match at that position. I then converted these possible base matches per position into degenerate IUPAC bases to form an estimated degenerate motif. The CEBPB (MA0466.2) calculated motif was VTKDYRHAAY, and the CTCF (MA0139.1) calculated motif was NNNMCDSNAGRDGDHRVNN.

### C.2. Parallelized searches increase speed as expected

Motif searches are independent, and therefore embarrassingly parallel, by contig and strand. This tool can take advantage of multiple processors for independent processing when they are available. For this test, I used the primary build of the GRCh38 human genome (Ensembl 91). I benchmarked the performance for multiple processor usage by searching both motifs in the human reference FASTA (sense and antisense) for ten iterations each of using 1-10 processors. I performed the benchmarks on a server with an Intel i7-3930k 3.20 GHz processor and 32 GB of RAM running Ubuntu 16.04.1 64-bit and Python 2.7.12. As expected, the single-processor runs were the slowest, and the runtime decreased noticeably with additional processors (**Fig. 1**). It is worth noting that the advantage of using multiple cores in this benchmark peaked at approximately 6 processors and stayed stable after that. The length and composition of the motifs is important, as demonstrated by different runtimes for each motif. Longer motifs, unsurprisingly, take longer to search compared to shorter motifs.

**Fig. 1.**
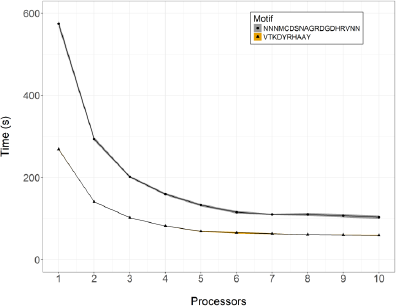
Runtimes for variable processor usage. Above are the runtimes for two motifs on the same system using 1-10 processors. The dots represent the means and the ribbons show the standard deviation for ten iterations of each condition.

## D. Summary

The lack of portable, general-purpose motif-finding tools for uses such as genome annotation is a significant barrier for the discovery of motifs in new / non-model genomes. The rapid increase in the number of available whole-genomes only amplifies this problem. Motif scraper aims to fill this gap. This tool has cross-platform compatibility and a permissive license for broad reuse. The runtime for annotation of relatively degenerate nucleotide sequences is fast, on the order of minutes for a whole-genome using multiple processors. The FASTA format allows for flexible input, ranging from whole genomes down cDNA sequences and plasmids. It could also be used to search for potential microRNA binding seed sequences in 3’ UTRs to predict potential partners for organisms not available in TargetScan (Agarwal, et al., 2015).

It is worth noting that the performance of parallel processing is highest with few relatively large contigs, i.e. reference genomes. The algorithm can be applied to smaller contigs, such as 3’ UTRs from a whole-genome to identify microRNA binding sites. However, the performance decreases appreciably with many short contigs. This limitation could be overcome by instead processing a batch of contigs per core to limit the number of data transfer operations. Overall, the broad operating system compatibility, use of a standard input format, and relative speed help support motif scraper as an important tool for non-model organisms and annotation of non-standard motifs.

## E. Acknowledgements

Special thanks to Dr. Karyn Meltz Steinberg for her helpful discussions during the development of this tool.

